# LncRNA YIYA enhances pancreatic cancer proliferation under high-glucose conditions through RAS–PKM2–mediated metabolic reprogramming that reinforces the Warburg phenotype

**DOI:** 10.1101/2025.10.30.685546

**Authors:** Nachiket Dattatray Garge, Ankita Sharma, Sudeshna Mukherjee, Rajdeep Chowdhury

## Abstract

Diabetes and the resulting hyperglycemia are metabolic complications frequently linked to pancreatic cancer (PC). In this study, we observe that the long non-coding RNA (lncRNA)-YIYA, which resides in a genomic region (1q32), frequently altered in various cancers, is overexpressed under conditions of chronic or intermittent high glucose (HG; 25 mM) compared to normal glucose (NG; 5.5 mM) in pancreatic ductal adenocarcinoma (PDAC) cells. It plays a pivotal role in regulating cellular metabolism-linked proliferation of PDAC cells, both under 2D and 3D conditions. Analysis of clinical datasets suggested that patients with elevated YIYA expression generally experience reduced survival in pancreatic cancer. Mechanistically, YIYA was found to regulate cellular energy metabolism and promote a Warburg phenotype, thereby facilitating the rapid proliferative status of PDAC cells. YIYA-mediated effects were found to be dependent on KRAS. Interestingly, the RIP assay revealed that YIYA interacts not only with KRAS but also with an important enzyme catalyzing the final step of the biochemical process, glycolysis, pyruvate kinase M2 (PKM2). YIYA was found to be predominantly localised in the cytoplasm, protecting the stability of the KRAS protein under a hyperglycaemic state. Importantly, YIYA knockdown enhanced autophagy-mediated degradation of KRAS. Our study is the first to identify YIYA as a glucose-responsive lncRNA in PDAC cells, and to reveal a novel YIYA-mediated connection between oncogenic KRAS signalling and metabolic reprogramming. Targeting YIYA represents a promising therapeutic strategy by disrupting the KRAS–PKM2–Warburg axis, thereby sensitizing PDAC cells.

**Graphical abstracts figure: Proposed model illustrating the effect of hyperglycemia on lncRNA YIYA–mediated regulation of KRAS signaling and metabolic reprogramming in pancreatic cancer:** Under hyperglycemic conditions, elevated glucose uptake through GLUT (Glucose Transporter) enhances glycolytic flux, leading to increased levels of PEP (Phosphoenolpyruvate) and activation of PKM2 (Pyruvate Kinase M2), which results in increased pyruvate flux toward the LDH (Lactate Dehydrogenase) pathway and subsequent increased lactate production, promoting the Warburg effect. Downregulation of MPC (Mitochondrial Pyruvate Carrier) also supports reduced shuttling of pyruvate towards the TCA (Tricarboxylic Acid) cycle. On the other hand, constitutively activated master regulator KRAS activates the downstream proliferative signalling via AKT and other DTs (Downstream targets), which upregulates various genes, including lncRNA YIYA aka LINC00538. YIYA exists in a functional complex with KRAS and PKM2, further reinforcing glycolytic and proliferative signalling. Additionally, YIYA suppresses autophagy-mediated degradation of KRAS. Together, these events contribute to hyperglycemia-driven epigenetic changes, metabolic reprogramming, and tumor progression in pancreatic cancer cells.

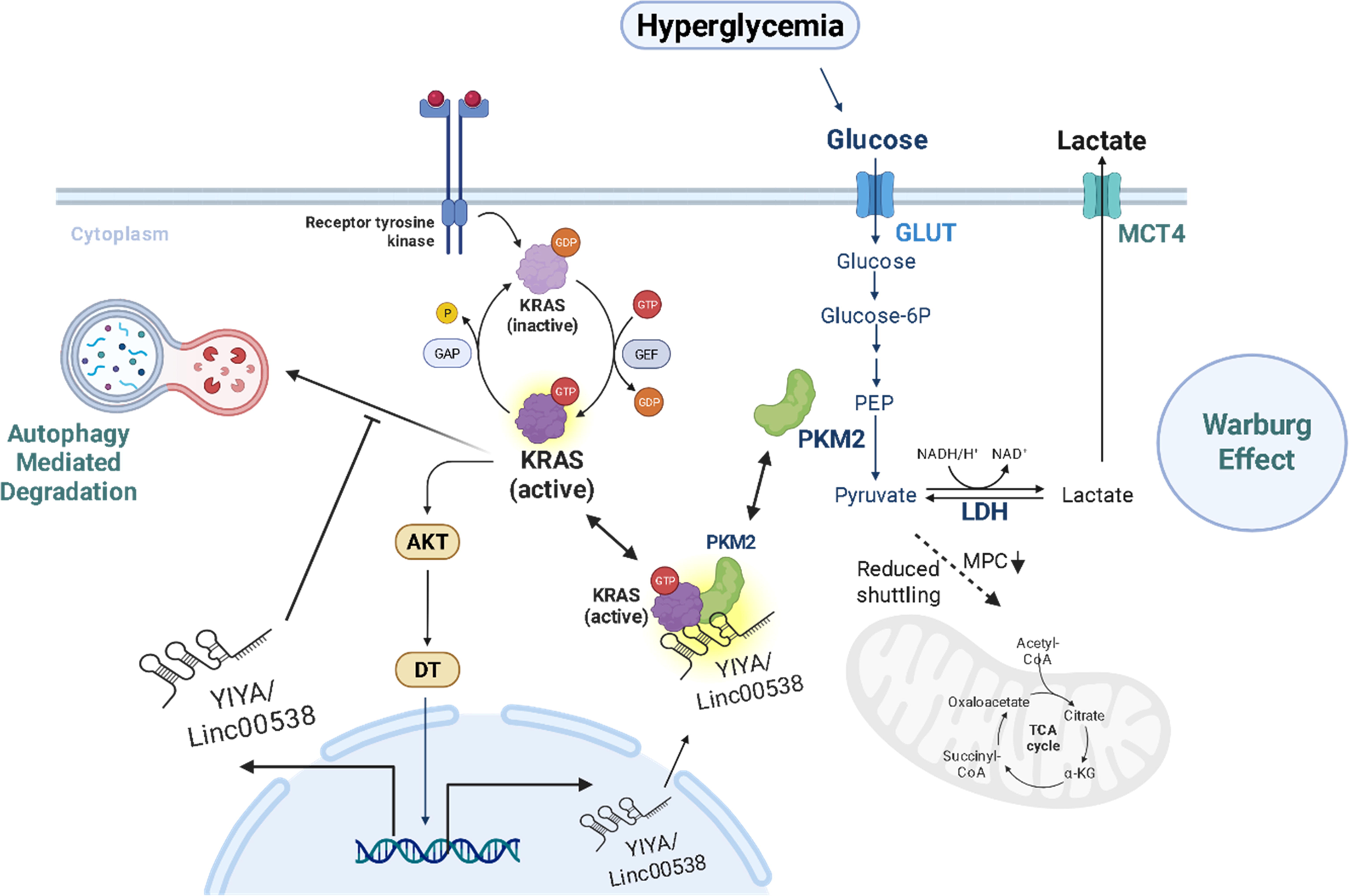

## Introduction

While our understanding of cancer and treatment methods has advanced significantly, pancreatic cancer continues to be among the most lethal types of malignancies worldwide. Although considerable progress has been made in understanding cancer biology and developing novel therapies, pancreatic cancer (PC) continues to rank among the most lethal malignancies worldwide. The overall five-year survival rate is below 10%. Among pancreatic tumors, pancreatic ductal adenocarcinoma (PDAC) represents more than 90% cases. Due to its typically late diagnosis, often at severe or metastatic stages, patients are left with very limited therapeutic options. (1). PDAC exhibits strong chemoresistance, a densely fibrotic and desmoplastic tumor microenvironment (TME), and an aggressive tumor biology. These combined features contribute to the disease’s poor prognosis and difficulty in developing effective treatments (2).

In addition to its aggressive pathology, PDAC profoundly affects whole-body metabolism. One well-documented feature is its strong link with diabetes and impaired glucose control (3). More than half of individuals diagnosed with PDAC show signs of hyperglycemia or develop new-onset, pancreatogenic (Type 3c) diabetes even before the tumor is detected. This form of diabetes can both contribute to and result from pancreatic cancer, often emerging from pancreatic injury due to chronic inflammation or the tumor itself. Elevated blood glucose, therefore, lies at the center of this two-way relationship between metabolic dysfunction and PDAC progression (4). However, one of the important risk factors of PDAC that occupies the hub of this bi-directional relationship is elevated blood glucose levels, also known as hyperglycemia.

Elevated blood glucose or hyperglycemia is thought to be both a result and a contributing factor in pancreatic tumor progression. High-glucose (HG) conditions can promote oncogenic transformation by altering metabolic pathways, increasing nutrient absorption, and activating glucose-dependent signaling (5). A well-recognized feature of this metabolic rewiring is the Warburg effect, where cancer cells rely on aerobic glycolysis and produce lactate despite sufficient oxygen availability (6). This metabolic strategy enables rapid cell growth by providing precursors required for synthesizing nucleotides, amino acids, and lipids. High-glucose conditions are known to drive the Warburg phenotype by stimulating the expression of glucose transporters, such as GLUT1, and crucial glycolytic enzymes, including HK2 and PKM2. These changes collectively boost glycolytic flux and strengthen the malignant metabolic program (7). Among these enzymes, PKM2 holds a central role in cancer metabolism. Unlike the constitutively active PKM1 found in healthy tissues, PKM2 functions as an allosterically regulated isoform capable of shifting between active tetrameric and less active dimeric states. This adaptability allows tumor cells to redirect glycolytic intermediates toward biosynthetic pathways, supporting continuous growth and disease progression (8).

Herein, KRAS is one of the important oncogenes that regulates these glycolytic metabolic adaptations in PC. KRAS is mutated in over 90% of PDAC cases (9). Oncogenic KRAS is essential for metabolic reprogramming as well as the activation of the traditional MAPK and PI3K signaling pathways. KRAS promotes the uptake of glucose, increases the activity of glycolytic enzymes, and facilitates the flow of metabolic intermediates into antioxidant and biosynthetic pathways. These pathways include the biosynthesis of hexosamine and the pentose phosphate pathway (10). Nevertheless, little is known about the molecular regulatory systems that govern glucose metabolism and KRAS signaling in PC during hyperglycemia.

Importantly, epigenetic remodelling is one of those regulatory systems that involves changes in gene expression that occur without alterations in the DNA sequence. In this regard, hyperglycemia is now known to regulate epigenetic remodelling by altering DNA methylation patterns, histone modifications, and chromatin status (open or closed chromatin) (11,12). These changes are often enzyme-driven. Interestingly, increased glucose levels can have an impact on the concentration of main metabolites such as acetyl-CoA, S-adenosylmethionine (SAM), and α-ketoglutarate. These metabolites act as substrates or cofactors for enzymes like histone acetyltransferases (HATs), DNA methyltransferases (DNMTs), and histone demethylases (KDMs), respectively. This shows a strong link between hyperglycemia and epigenetic regulation. (12). For example, high glucose has been shown to increase histone acetylation *via* increased acetyl-CoA. This leads to an open chromatin structure, and in turn, transcriptional activation of oncogenes and metabolic regulators (13). Such dynamic and reversible changes provide a mechanism through which metabolic states can influence gene expression.

Long non-coding RNAs (lncRNAs) are now known to be key regulatory molecules in the framework of epigenetics. These transcripts, longer than 200 nucleotides and lacking coding potential, can modulate gene expression at transcriptional, post-transcriptional, and chromatin levels (14). Importantly, many lncRNAs are regulated by metabolic changes. For example, PVT1, located near the MYC oncogene, has been shown to regulate glucose metabolism and redox homeostasis in PDAC (15). Similarly, MALAT1 has been implicated in regulating glycolysis and autophagy *via* modulation of the mTOR signalling (16). Other lncRNAs like HULC, LINC01133, and H19 have also been associated with glucose-responsive signalling and metabolic reprogramming in multiple cancers. Despite these advances, the role of lncRNAs as mediators of hyperglycemia-induced epigenetic and metabolic rewiring in PDAC remains poorly understood.

In the current study, we analysed the consequences of high glucose (25 mM) on lncRNA-mediated metabolic regulation in PDAC cell lines. Our results show that elevated glucose levels induce the expression of the novel long non-coding RNA YIYA, a transcript previously reported in metabolic regulation of breast cancer (17), but not yet studied in the context of PDAC. YIYA, which resides in a genomic region (1q32) frequently altered in various cancers (18). Here, we identify a novel interaction between YIYA and KRAS, implicating this lncRNA in the modulation of both metabolic and oncogenic signalling pathways. Notably, we demonstrate that YIYA stabilizes the proteins-KRAS and pyruvate kinase 2 (PKM2) by preventing their autophagy-mediated degradation, thus facilitating proliferation. Collectively, our findings reveal a previously unrecognized KRAS–YIYA–PKM2 axis, activated under hyperglycaemic conditions, that promotes PDAC progression through the coordinated regulation of oncogenic and metabolic networks.

## Materials and Methods

### Reagents and Chemicals

Propidium iodide (PI; #P4864), TRI reagent® (#T9424), Bradford Reagent (#B6916), Chloroquine (CQ, #C6628), and RIPA Buffer (#R0278) were obtained from Sigma. MTT (#33,611) was manufactured by SRL. SYBR Green Supermix (#1725121), Clarity™ Western ECL Substrate (ECL, # 1705061), and cDNA synthesis kit (#1708891) were procured from Biorad. Lipofectamine 3000 (#L3000-001), Annexin-V FITC-Conjugate (#A13199), and Annexin V binding buffer (#V13246) were procured from Invitrogen. Alexa FluorTM Plus 555 Anti-rabbit IgG antibody (#32732) was procured from Invitrogen. Glucose Uptake-Glo™ Assay (#J1341), and Lactate-Glo™ Assay (#J5021) were purchased via Promega. siYIYA was synthesized by Genecust, France. KRAS inhibitor (HY-130149), AKT inhibitor (HY-10115), Protein A/G magnetic beads (HY-K0202), and PKM2 inhibitor (HY-103617) were procured from MedChemExpress. Extracellular Oxygen Consumption Assay (ab197243) was obtained from AbCam.

### Bioinformatics Analysis

For bioinformatics analysis, survival probability plots for lncRNA YIYA were obtained from the lnc2cancer database (http://www.bio-bigdata.com/lnc2cancer), which integrates experimentally supported associations between lncRNAs and various cancers. Kaplan–Meier survival plots for KRAS were generated using the GEPIA2 platform (http://gepia2.cancer-pku.cn), which provides RNA-sequencing expression and survival data based on TCGA and GTEx datasets. Survival probability was analyzed using the log-rank test

### Cell Culture and Exposure to Glucose

MIA PaCa-2 and PANC-1 pancreatic cancer cells (NCCS, Pune) were cultured in DMEM (HiMedia) containing 10% FBS, 1% penicillin– streptomycin, and either 5.5 mM (NG) or 25 mM (HG) glucose. Cells were maintained at 37 °C with 5% CO□ and serum-starved for 12 hours once they reached 65–70% confluence. For intermittent high-glucose exposure, cells were seeded to reach 65%–70% confluence at the start and cultured in either normal glucose or high-glucose media. The intermittent regimen consisted of 12-hour cycles of NG → HG → NG (media exchanged every 12 h with a gentle PBS rinse to minimize carryover). RNA was harvested at 36 h and 72 h time points.

### Cell Viability Assay

Cell viability was assessed by the 3-(4,5-dimethylthiazol-2-yl)-2,5-diphenyltetrazolium bromide (MTT) assay. Following treatment, MTT reagent was added to each well, and cells were incubated for 4 hours at 37°C. After formation, the formazan crystals were dissolved in DMSO, and the absorbance was measured at 570 nm with a reference wavelength of 630 nm using a Multiskan Sky spectrophotometer (Thermo Fisher Scientific). The percentage of viable cells was then calculated using the equation below: % Cell Viability = (Average absorbance of treated cells / Average absorbance of control cells) × 100 (19).

### Crystal Violet Staining

The cells were seeded in a 6-well plate at a density of 1 × 10^6^ cells/well. The cells were treated as per the experimental design. After the treatment period, cells were washed three times with 1X PBS and fixed with ice-cold methanol at –20°C for 20 minutes. Following fixation, cells were again washed three times with PBS and incubated with 0.5% crystal violet solution for 10 minutes at room temperature. Excess stain was gently removed, plates were air-dried, and images were captured for analysis.

### Wound Healing Assay

The cells were seeded in a 6-well plate at a density of 1 × 10^6^ cells/well. A linear scratch was made at the centre of each well using a sterile 10 µL pipette tip. After scratching, cells were washed to remove debris and incubated in defined treatment conditions. Images of the wound area were captured at 0 and 48 hours using a Zeiss Primovert microscope.

### RNA Extraction, cDNA Synthesis and qRT-PCR

Total RNA was isolated using TRI reagent according to the manufacturer’s instructions. cDNA was synthesized using the iScript cDNA synthesis kit (Bio-Rad). Quantitative PCR was performed on the QuantStudio 3 system (Applied Biosystems) using gene-specific primers and SYBR Green dye for detection. GAPDH served as the internal reference gene, and relative transcript levels were quantified using the 2^ΔΔCt method as described by Livak and Schmittgen (19).

### Western Blotting

Cells were cultured in 10 cm dishes and harvested at designated time points. Protein lysates were extracted using RIPA buffer (containing protease inhibitor), and the total protein concentration was determined using the Bradford assay, with absorbance measured at 595 nm on a Multiskan Sky spectrophotometer (Thermo Fisher Scientific). Lysates were mixed with 5X loading dye and boiled at 100°C for 10 minutes. Proteins were separated by SDS-PAGE and transferred onto polyvinylidene fluoride (PVDF) membranes. 3% BSA was used to block the membranes. Membranes were probed overnight at 4 °C with primary antibodies, stripped when necessary, and re-probed for additional targets. Bands were visualized by ECL. The ChemiDoc machine was used and measure with ImageJ relative to control protein levels (20).

### Annexin V/Propidium Iodide (PI) Apoptosis Assay

Approximately 6 × 10[ cells were seeded per dish. Upon reaching 70%–80% confluency, cells were subjected to the indicated treatments. Post-treatment, cells were trypsinized, washed with PBS, and resuspended in 500 µL of 1X binding buffer (BD Biosciences). To assess cell death, 3 µL each of Annexin V and propidium iodide (PI) was added to the cell suspension and incubated in the dark at room temperature for 30 minutes. Samples were acquired using a Beckman Coulter CytoFLEX flow cytometer, and data were analyzed using CytExpert software. The two left quadrants of the flow cytometry plots represent the percentages of early and late apoptotic cells. The fold change in percentage of cell death was quantified and represented as a bar graph (20).

### Cell Cycle Analysis by Propidium Iodide Staining

To determine the distribution of cells across different cell cycle stages, cells were plated in 6-cm dishes. Following that, the cells were transfected with siRNA of YIYA. Post-transfection, cells were collected, washed with PBS, and centrifuged at 2500 rpm for 10 minutes at 4 °C. For fixation, chilled 70% ethanol was added dropwise to the cell suspension with gentle mixing, and the samples were kept overnight at 4 °C. Following that, the cells were rinsed with PBS and incubated in 500 µL of PBS supplemented with 3 µL of propidium iodide for 10 minutes at room temperature, protected from light. The measurement of DNA content was performed using a CytoFLEX flow cytometer, and to quantify the proportions of cells in each phase of the cell cycle, CytExpert software was utilized.

### Transient Transfection & Knockdown

Knockdown of YIYA was performed using custom-designed siRNAs. The sequence of the siRNA is as follows: 5’-GGUACUAAGAUAUCUGGAAAUCAGC-3’. Approximately 6 × 10[ cells were seeded per dish. The following day, cells were transfected with 40 nM of the respective siRNAs using Lipofectamine 3000 reagent (Thermo Fisher Scientific) according to the manufacturer’s instructions. A scrambled siRNA was transfected in parallel as a negative control. After the experimental time point, cells were then harvested for RNA extraction and subjected to downstream analyses, including assessment of cell death (21).

### Immunofluorescence Microscopy

Cells (1.5 × 10[) were seeded on coverslips, treated as indicated, and then the cells were treated with 2% paraformaldehyde for 15 minutes for the fixation process. After permeabilization with 0.2% detergent (Triton X-100), samples were blocked with 1% BSA. Following that, primary antibodies (1:1000) incubation was done overnight at 4 °C. Then, samples were incubated with FITC-or TRITC-conjugated secondary antibodies (1:5000) for 90 minutes in the dark. DAPI antifade medium was used to mount coverslips. Images were taken with a Zeiss confocal microscope. Image processing was performed using Zen 2.3 SP1 software.

### Extracellular Oxygen Consumption Rate (OCR) Assay

To evaluate extracellular oxygen consumption, the Abcam kit (ab197243) was used according to the manufacturer’s recommendations. Approximately 5,000 cells were added to a 96-well plate. Post treatment, the oxygen consumption reagent was introduced, and the wells were overlaid with high-sensitivity mineral oil to prevent oxygen diffusion from the air. Fluorescence was recorded for 4 hours using a Victor microplate reader. The dye was excited at 360–380 nm (maximum 380 nm), and emission was measured at 630–680 nm (maximum 650 nm).

### ATP Measurement

The CellTiter-Glo Luminescent Cell Viability Assay (Promega) was used to quantify metabolically active cells. Briefly, 50 µL reconstituted reagent and 50 µL cell suspension were mixed in each well. The cell lysis was induced by a brief shaking of the plate, followed by a 10-minute incubation at room temperature that allowed the stabilization of the signal. ATP levels, indicative of viable cells, were measured as relative light units (RLUs) using the Perkin Elmer VICTOR Multilabel Counter.

### RNA Fractionation

The cell fractionation protocol was adapted and modified from previously published methods. Cells were cultured to ∼80% confluency, trypsinized, and counted. For lysis, cells were resuspended in lysis buffer (50 mM Tris-Cl, pH 8.0, 100 mM NaCl, 0.5% NP-40, and 5 mM MgCl[) and centrifuged at 2,500 × g for 5 minutes at room temperature in 1.5 mL tubes. The supernatant was discarded, and the pellet was gently resuspended in 300 µL of ice-cold 0.25X lysis buffer with intermittent shaking. This was followed by centrifugation at 13,000 × g for 2 minutes at 4°C. The supernatant, representing the cytoplasmic fraction, was carefully transferred to a fresh 1.5 mL tube, while the remaining pellet was retained as the nuclear fraction. To each fraction, Buffer RLT (Qiagen) supplemented with 1% β-mercaptoethanol (10 µL per 1 mL) was added: 1050 µL for the cytoplasmic and 600 µL for the nuclear fraction. Ethanol was then added to each: 750 µL of 90% ethanol to the cytoplasmic and 600 µL of 70% ethanol to the nuclear lysate. Samples were transferred to RNeasy MinElute spin columns (Qiagen), placed in 2 mL collection tubes, and centrifuged at 8,000 × g for 15 seconds. Flow-through was discarded, and columns were washed sequentially with 700 µL and 300 µL of Buffer RW1, each followed by centrifugation at 8,000 × g for 15 seconds. To dry the membrane, spin columns were placed in fresh 2 mL collection tubes and centrifuged at maximum speed for 5 minutes with the cap open. Finally, columns were placed in new 1.5 mL collection tubes, and RNA was eluted by adding 30 µL of RNase-free water directly onto the membrane. The columns were centrifuged at full speed for 1 minute to collect the RNA. The integrity of fractionation was confirmed using GAPDH and NEAT1-2 as a cytoplasmic and nuclear control, respectively. (18).

### Glucose uptake assay

Glucose uptake was measured using the Glucose Uptake-Glo™ Assay Kit (Promega, Cat. #J1341) following the manufacturer’s instructions. Cells grown in high-glucose conditions were transfected with siYIYA or control siRNA for knockdown studies. On the assay day, the medium was replaced with glucose-free PBS, and cells were treated with 1 mM 2-deoxyglucose for 10 minutes at room temperature. Sequential addition of Stop and Neutralization buffers (25 µL each) stopped the uptake, 100 µL of 2DG6P detection reagent followed. Post incubation for 30–60 minutes in the dark, luminescent signals were measured using a PerkinElmer VICTOR® Nivo plate reader. No-2DG and Stop-before-2DG controls were included for background correction. A 2DG6P standard curve (0–30 µM) was used to quantify uptake, and values were normalized to cell number.

### Lactate Quantification

Extracellular lactate levels were quantified using the Lactate-Glo™ Assay Kit (Promega, Cat. #J5021) according to the manufacturer’s instructions. Briefly, culture media were collected from cells grown in normal glucose (5.5 mM) or high glucose (25 mM) DMEM and diluted 1:40 in PBS. Fifty microliters of each diluted sample were incubated with an equal volume of freshly prepared Lactate Detection Reagent in white opaque 96-well plates. After 60 minutes of incubation at room temperature in the dark, luminescence was recorded using a PerkinElmer VICTOR® Nivo multimode plate reader. A standard curve (0–200 µM) was generated using known lactate concentrations, and sample concentrations were calculated by linear regression after background subtraction (only media wells were used for background subtraction).

### Co-immunoprecipitation (Co-IP) Assay

Cells were cultured in 10 cm dishes and harvested at designated time points. Protein lysates were extracted using RIPA buffer (containing protease inhibitor), and the total protein concentration was determined using the Bradford assay, with absorbance measured at 595 nm on a Multiskan Sky spectrophotometer (Thermo Fisher Scientific). Total protein (700 µg) was used for pulldown with a specific antibody. The antibody was incubated with Protein A/G magnetic beads before the pulldown experiment. The pulled-down lysate was added to 70 µL Lameli buffer. Proteins were separated by SDS-PAGE and transferred onto polyvinylidene fluoride (PVDF) membranes. 3% BSA was used to block the membranes. Membranes were probed overnight at 4°C with the primary antibodies of interest. Bands were visualized by ECL. The ChemiDoc machine was used and measure with ImageJ relative to control protein levels (20).

### RNA Immunoprecipitation (RIP) Assay

The RIP assay was employed using the Magna RIP™ RNA-Binding Protein Immunoprecipitation Kit (Merck, #17-700). MIA PaCa-2 cells were lysed in RIP lysis buffer supplemented with RNase and protease inhibitors. Magnetic Protein A/G beads were incubated with 3 µg of the specific antibody or normal IgG (control) at room temperature for 30 min. The cleared lysate was incubated with the antibody-bead complex at 4 °C for 3 hours with rotation. After washing with RIP wash buffer, the RNA–protein complexes were digested with proteinase K. The RNA was then extracted using a phenol:chloroform:isoamyl alcohol solution, followed by ethanol precipitation. Purified RNA was resuspended in RNase-free water and used for downstream analysis by RT-qPCR.

### Chromatin Immunoprecipitation (ChIP)

The MAGnifyTM Chromatin Immunoprecipitation System (Thermo Fisher Scientific, CA, USA) was used to perform chromatin immunoprecipitation (ChIP) experiments according to the manufacturer’s protocol. In summary, cell pellets were obtained after treatment and then reconstituted in PBS. Crosslinking was done for 10 min using 1% (v/v) formaldehyde. Thereafter, 0.125 M glycine was added to the cells to stop the cross-linking reaction. Cells were washed with PBS containing protease inhibitors before being lysed in a kit-provided SDS lysis solution. After being sonicated for 30 seconds with chromatin, the lysate was allowed to cool over ice for 30 seconds multiple times to ac5hieve a chromatin size of 0.3–0.5 kb. Lysates were then centrifuged, and the supernatant was incubated for 2 h with H3K4me3 and H3K27me3 antibodies. Before the antibody was added, input samples were obtained from the crude lysates, and they were kept at 20°C until extraction. Protein-DNA complexes were eluted after antibody incubation, and then the protein-DNA complexes were reversed by the cross-linking buffer. DNA was purified by utilizing the kit’s magnetic beads. Using promoter primers, qPCR with SYBR reagent was performed for the input DNA samples and ChIP-enriched samples in a QuantStudio 3 qRT-PCR system. For the ChIP experiment, an anti-IgG antibody was used as a negative control.

### Spheroid Assay

MIA PaCa2 cells were cultured as 3D spheroids in 96-well round-bottom plates by seeding 3,000 cells per well in complete DMEM supplemented with 0.2% methylcellulose to enhance spheroid formation. Spheroids were incubated at 37°C with 5% CO[. Uniform spheroids typically formed within 1-2 days. Media was gently replaced every 2 days to preserve spheroid integrity. Treatments were administered by partial media replacement, and spheroids were processed for RNA/protein extraction or bright-field imaging.

### Statistical Analysis

Data are expressed as mean ± SD from at least three independent experiments. Statistical analyses were carried out using GraphPad Prism 8 (Windows). Depending on the experimental setup, significance between groups was assessed using Student’s t-test or one-/two-way ANOVA, followed by appropriate post hoc tests (Bonferroni, Tukey’s, Dunnett’s, or Sidak’s). A p-value < 0.05 was considered statistically significant. Statistically significant differences are indicated by symbols * as denoted in the figure legends.

## Results

### High-glucose exposure metabolically drives PDAC cell proliferation and induces lncRNA YIYA expression in both 2D and 3D culture conditions

Previous work from our lab has demonstrated that sustained high glucose (HG) conditions promote pancreatic ductal adenocarcinoma (PDAC) cell proliferation and survival through upregulation of genes, including proliferating cell nuclear antigen (PCNA) and Ki67 (22). While these findings provide strong evidence for the link between chronic hyperglycemia and tumour growth and aggressiveness, we wanted to explore further whether HG-mediated effects are dynamic or stable in nature. To evaluate the same, we switched the cells growing in NG to HG and *vice versa*. MIA PaCa-2 cells, initially adapted to either normal glucose (NG, 5.5 mM) or high glucose (HG, 25 mM) conditions, were switched to reverse glucose concentration, and the expression of Ki67 and PCNA was monitored. The transcript and protein analysis revealed an increased expression of proliferation markers when cells were switched from NG to HG (**Fig. 1a-c**) and contrasting decreased levels under the *vice versa* condition (**Fig. 1d-f**). Simultaneously, immunofluorescence staining of PCNA revealed a marked increase in nuclear localization of PCNA in HG conditions in the switch experiments (NG to HG and HG to NG) **(Supplementary Fig. 1a-b**).

**Figure 1:**
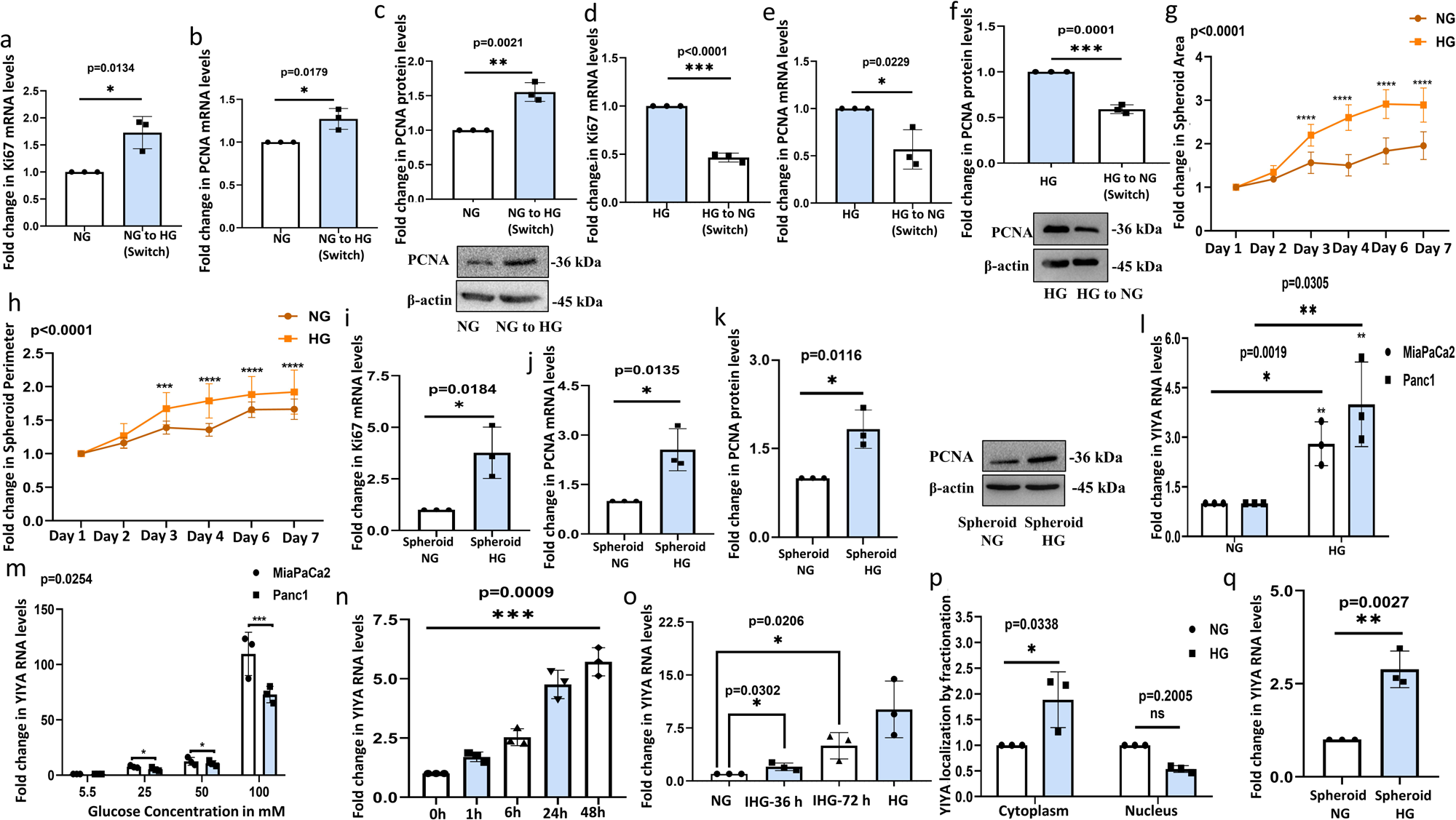
PDAC cells show increased proliferative ability and associated upregulation of lncRNA YIYA under high glucose in 2D and 3D culture conditions. **(a)** Relative mRNA expression of Ki67 and **(b)** PCNA, and **(c)** protein expression of PCNA in MIA PaCa-2 cells cultured under NG or switched from NG to HG conditions. **(d)** Relative mRNA expression of Ki67 and **(e)** PCNA, and **(f)** protein expression of PCNA in MIA PaCa-2 cells cultured under HG or switched to HG to NG conditions. **(g)** Change in area and **(h)** perimeter of spheroids of MIA PaCa-2 cells grown in NG or HG conditions, from day 1 to day 7. **(i)** Relative mRNA expression of Ki67 and **(j)** PCNA, and **(k)** protein expression of PCNA in spheroids of MIA PaCa-2 cells cultured under NG or HG conditions. **(l)** Relative expression of lncRNA YIYA in MIA PaCa-2 and PANC-1 cells cultured under NG or HG conditions. **(m)** Relative expression of lncRNA YIYA in MIA PaCa-2 and PANC-1 cells cultured under NG or HG conditions in a dose and **(n)** time-dependent manner. **(o)** Relative expression of lncRNA YIYA in MIA PaCa-2 cells cultured under NG, HG, and intermittent high glucose (IHG) (NG→HG→NG cycles of 12 hr) conditions. **(p)** LncRNA YIYA localization in NG and HG conditions through RNA fractionation followed by qPCR. **(q)** Relative expression of lncRNA YIYA in spheroids of MIA PaCa-2 cells cultured under NG or HG conditions. The glucose concentration used for NG is 5.5 mM, and for HG is 25 mM. Unless otherwise specified, treatments were carried out for 48 h. For quantitative comparisons, bar graphs representing gene expression or protein levels were normalized to the respective housekeeping gene or loading control (either GAPDH, NEAT1-2, or β-actin). Fold change values shown in the graphs are expressed relative to the control, which was set to 1. All results are presented as mean ± SD from at least three independent experiments (n = 3). For comparisons involving multiple groups across varying concentrations and/or time points, two-way ANOVA followed by Tukey’s multiple comparisons post-test was performed. For pairwise comparisons between two groups, unpaired t-tests were applied. Statistical significance is indicated as (*) p < 0.05, (**) p<0.01, and (***) p < 0.001 versus the control.

A three-dimensional (3D) cell culture often reflects an optimal growth scenario for cells and mimics a more physiologically relevant tumour microenvironment (23). Going ahead, we therefore optimized a protocol for spheroid culture of the MIA PaCa-2 cells. Importantly, the cells formed larger spheroids under HG conditions compared to NG from day 3 onwards, as evident from the bright field images **(Supplementary Fig. 1c**) and quantification of spheroid area and perimeter (**Fig. 1g-h**). Importantly, in 3D conditions as well, mRNA expression of the propagation markers Ki-67 and PCNA was upregulated in HG conditions compared to NG (**Fig. 1i-j**). Immunoblotting further revealed elevated PCNA protein expression in HG compared to NG, when cells were grown in 3D spheroids, supporting an enhanced proliferation (**Fig. 1k**).

Tumour cells have been abundantly explored for inheriting mutations causative of stable phenotypic modifications. However, existing literature suggests an epigenetic bias towards transitory or dynamic alterations (24). In this regard, to understand the molecular dependency of PDAC cells to HG-induced proliferative adaptations, we explored the expression of the lncRNA YIYA. LncRNA YIYA expression was found to be consistently increased in both the PC cell lines studied, PANC-1 and MIA PaCa-2, under HG conditions (**Fig. 1l**). We further observed a dose-dependent increase in YIYA expression with escalating glucose concentrations in MIA PaCa-2 cells (**Fig. 1m**). Time-dependent expression kinetics further demonstrated an early induction of YIYA as early as 1 h post-HG exposure (**Fig. 1n**). Recent clinical and preclinical studies have highlighted that intermittent spikes in blood glucose, rather than sustained hyperglycemia alone, might have unique biological effects on tumor progression and cellular stress responses. Such intermittent high glucose (IHG) patterns are increasingly recognized in diabetic patients and have been associated with metabolic reprogramming in cancer cells (25,26). Moving forward, we therefore wanted to examine how IHG exposure influences the expression of lncRNA YIYA in PDAC cells. Glucose switch after every 12 h from NG to HG, and the reverse was continued for specific time periods (36 h and 72 h), and RNA was extracted. Such intermittent spikes of high glucose also showed a rapid induction of YIYA mRNA; however, the expression was comparatively lower than in cells under chronic HG (**Fig. 1o**). An earlier report, on colorectal cancer, indicates that YIYA can interact with the catalytic subunit of the chromatin repressive complex-EZH2, and therefore, can be found in the nucleus, while studies on breast cancer by Zheng Xing et al (2018) indicated a contrasting cytoplasmic distribution of the lncRNA as well (17,27). To confirm its localization, we hence performed a subcellular fractionation. Importantly, YIYA was found to be predominantly localized in the cytoplasm under HG, suggesting a putative non-nuclear function in the current context (**Fig. 1p**). Importantly, YIYA expression was significantly upregulated in HG spheroids as well compared to NG, which clearly indicates that YIYA induction is conserved across both 2D and 3D culture conditions (**Fig. 1q**). The results from both 2D and 3D culture experiments are indicative of a significant upregulation of lncRNA YIYA under HG in PDAC cells. Importantly, while a switch from NG to HG resulted in a spike in YIYA expression, however, the reverse was not true. This might be indicative of a glucose-driven memory driving YIYA expression **(Supplementary Fig. 1d-e**).

### lncRNA YIYA is critical for sustaining PDAC cell proliferation, and its elevated expression correlates with reduced overall survival in pancreatic cancer patients

To investigate the functional significance of YIYA in PDAC progression, we performed siRNA-mediated knockdown of YIYA in MIA PaCa-2 and PANC-1 cells cultured under HG conditions. Trypan blue exclusion assay confirmed an increase in the population of non-viable cells in the siYIYA group (**Fig. 2a**). Consistently, crystal violet staining and phase-contrast imaging also revealed a visible decline in cell viability upon YIYA knockdown (**Fig. 2b and Supplementary Fig. 2a**). At the molecular level, transcriptional and protein analyses in both cell lines revealed reduced expression of Ki67 and PCNA following YIYA knockdown (**Fig. 2c-e**), which aligns with the reduced proliferative phenotype. Furthermore, Annexin V-PI staining confirmed an increase in the population of apoptotic cells in the siYIYA group, and in parallel, a significant elevation in caspase-3 transcript levels was observed upon YIYA knockdown (**Fig. 2f and Supplementary Fig. 2b**), supporting the induction of apoptotic signalling. We next examined how YIYA knockdown impacts cell cycle progression. An arrest in the G0/G1 phase and a concomitant reduction in S-phase population in MIA PaCa2 cells were shown in flow cytometry-based cell cycle analysis (**Fig. 2g**), suggesting a blockade in cell cycle progression. Moreover, YIYA-depleted cells showed markedly reduced wound healing potential at 48 hours under HG conditions, which hints towards impaired cell migration and/or proliferation **(Supplementary Fig. 2c**). To know the underlying epigenetic changes, we performed ChIP-qPCR targeting the PCNA upstream promoter region. Interestingly, ChIP-qPCR (**Fig. 2h**) analyses revealed decreased enrichment of H3K4me3, an open chromatin mark, and an increased enrichment of H3K27me3, a closed chromatin mark, at the PCNA promoter in the absence of YIYA (**Fig. 2h**), which suggests a YIYA-dependent indirect epigenetic regulation of the above genes. Given that a vast majority of PC patients harbour a pre-existing hyperglycaemic condition or often develop diabetes post-occurrence of PC (28), we wanted to explore the clinical relevance of YIYA expression in PC patients. A Kaplan–Meier survival analysis using data from the lnc2cancer database (http://www.bio-bigdata.net/lnc2cancer) revealed that patient survival in PC was inversely correlated with high expression of YIYA **(Supplementary Fig. 2d**).

**Figure 2:**
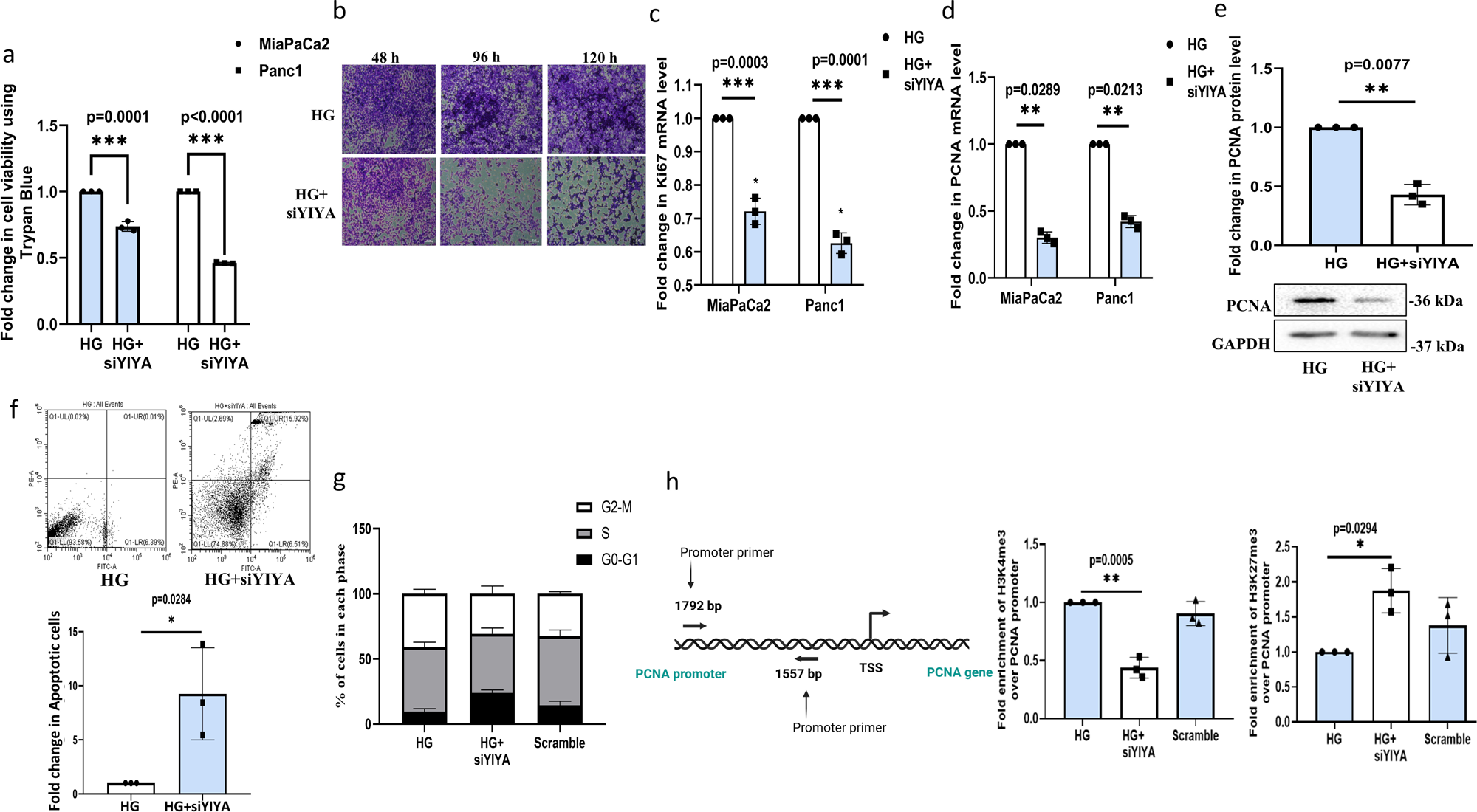
YIYA is essential for the proliferation of PDAC cells, and its expression is positively correlated with poor survival of PC patients. **(a)** Relative fold change in cell viability upon siRNA-mediated YIYA knockdown under HG conditions using trypan blue in MIA PaCa-2 and PANC-1 cells. **(b)** Crystal violet staining after incubation of MIA PaCa-2 cells in HG plus siYIYA (siRNA-mediated YIYA knockdown), as compared to the HG condition (Scale Bar: 50 µm). **(c)** Relative mRNA expression of Ki67 and **(d)** PCNA, and **(e)** PCNA protein expression upon siRNA-mediated YIYA knockdown under HG conditions in MIA PaCa-2 and PANC-1 cell lines. **(f)** Relative fold change in apoptotic cells cultured under HG and siRNA-mediated YIYA knockdown in MIA PaCa-2 cells. **(g)** Cell cycle analysis through flow cytometry post-YIYA knockdown in MIA PaCa-2 cells. **(h)** Enrichment of H3K4me3 and H3K27me3 at PCNA promoter under HG and post-YIYA knockdown, as analysed using ChIP-qPCR (Promoter primers were designed to target the PCNA promoter region, located 1.7 kb upstream of the transcription start site [TSS]). Input served as control. ChIP-qPCR illustration prepared with Biorender software. The glucose concentration used for NG is 5.5 mM, and for HG is 25 mM. The final concentration of siRNA used for transfection was 40 nM. Unless otherwise specified, treatments were carried out for 48 h. For quantitative comparisons, bar graphs representing gene expression or protein levels were normalized to the respective housekeeping gene or loading control (either GAPDH or β-actin). Fold change values shown in the graphs are expressed relative to the control, which was set to 1. All results are presented as mean ± SD from at least three independent experiments (n = 3). For comparisons involving multiple groups across varying concentrations and/or time points, two-way ANOVA followed by Tukey’s multiple comparisons post-test was performed. Statistical significance is indicated as (*) p < 0.05, (**) p<0.01, and (***) p < 0.001 versus the control. For pairwise comparisons between two groups, unpaired t-tests were applied.

### HG activates KRAS-dependent signalling that upregulates YIYA highlighting a convergence between oncogenic KRAS activity and metabolic adaptation in PDAC cells

KRAS is a central oncogenic driver in PC, playing a critical role in cell growth, survival, and metabolic regulation. It is now well established that PC patients harbouring mutations in RAS family members, including G12C and G12D, exhibit a significantly worse overall survival (29). This clinical association motivated us to further investigate the role of KRAS, if any, in our experimental models. Given its prevalence and impact, we sought to explore whether KRAS might influence YIYA expression under hyperglycaemic conditions. To understand the clinical significance of RAS pathway activation in PDAC, we referred to published studies that analysed patient data (30,31). As shown in **Fig. 3a**, patients harbouring higher expression of KRAS exhibited significantly worse overall survival. Importantly, the MIA PaCa-2 and PANC-1 cells used for this study harbour a KRAS G12C mutation and a heterozygous G12D mutation, respectively (32). These associations further motivated us to investigate the connection between hyperglycemia and KRAS signalling in our experimental model. We initially analysed the mRNA and protein expression levels of KRAS and its most prominent downstream partner, AKT, upon HG. Interestingly, qPCR and immunoblot analysis revealed increased expression of KRAS in both 2D and 3D culture conditions (**Fig. 3b-e**). Immunofluorescence analysis further validated increased KRAS expression in HG-treated cells (**Fig. 3f**). Consistently, AKT, a key downstream effector of KRAS, also showed elevated phosphorylation (pAKT; Thr308) (33), indicative of its active state under HG (**Fig. 3g**). To further investigate the KRAS dependency, we treated the MIA PaCa-2 cells with a KRAS inhibitor (Adagrasib) under HG conditions. In this regard, Adagrasib, known to dampen KRAS protein activity by binding to the KRAS G12C mutant (34,35), showed a marked diminishment in expression of KRAS and the downstream effector-ERK **(Supplementary Fig. 3a and b**). Importantly, this was associated with a significantly reduced expression of YIYA upon KRAS inhibition in MIA PaCa-2 cells cultured under 2D (**Fig. 3h**) and 3D (**Fig. 3i**) conditions as well. Importantly, in accordance with the proliferation-promoting role of YIYA, inhibition of KRAS also led to a drastic reduction in spheroid dimension and area when the cells were cultured under HG in 3D **(Supplementary Fig. 3c and d**). These results clearly indicate that YIYA expression is KRAS-dependent in a hyperglycaemic context. To further dissect this axis and to investigate whether the downstream AKT signalling plays a role in regulating YIYA, we pharmacologically inhibited AKT using a specific AKT inhibitor (PI103) (36). Successful inhibition of the pathway was confirmed by a steep downregulation in pAKT levels, shown by western blot image **(Supplementary Fig. 3e**). Interestingly, suppression of AKT signalling also led to a noticeable decrease in lncRNA YIYA expression (**Fig. 3j**), which indicates that YIYA may be downstream of the KRAS-AKT pathway and potentially regulated by its activity under hyperglycaemic conditions.

**Figure 3:**
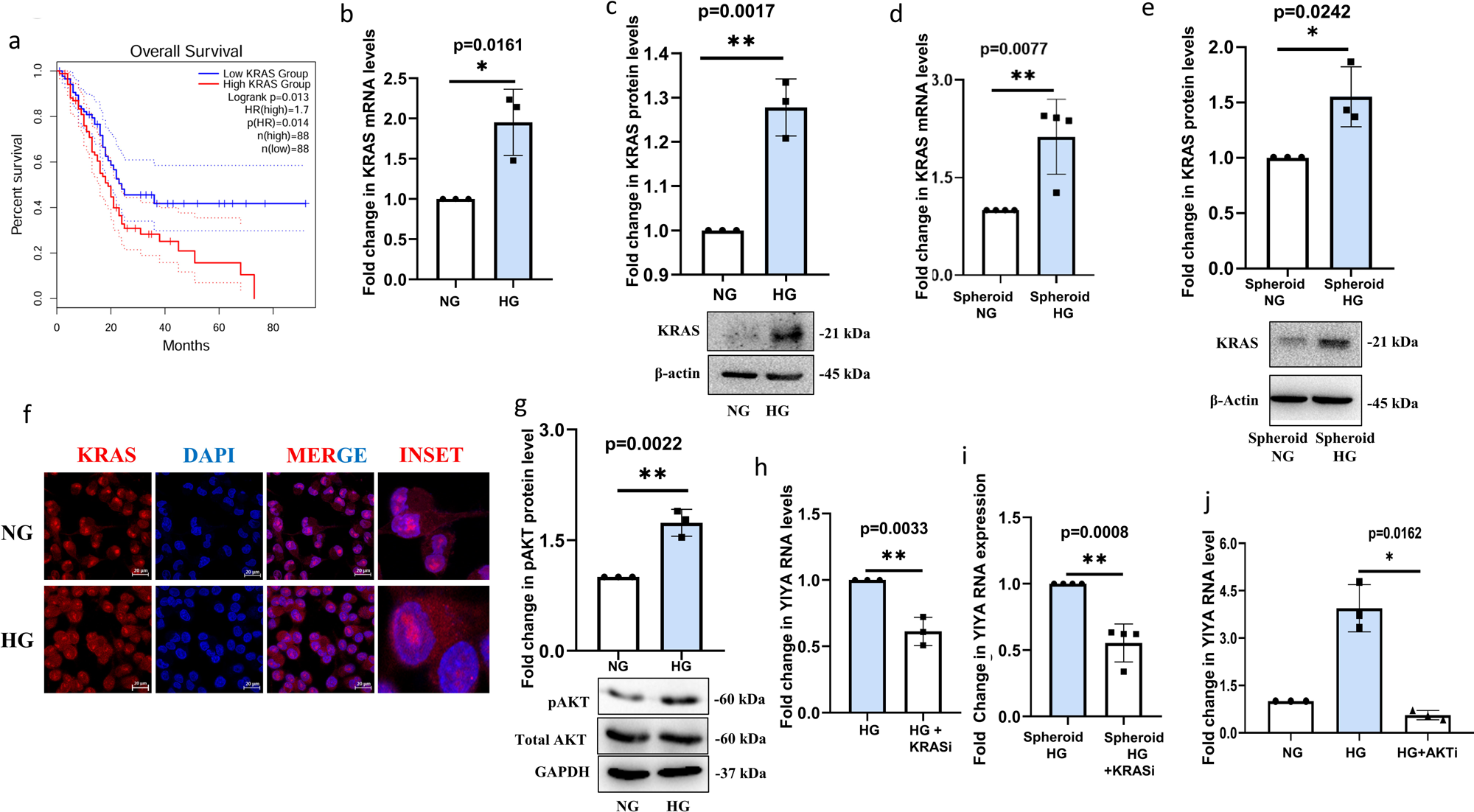
YIYA expression is regulated by KRAS signalling under HG conditions. **(a)** Kaplan–Meier survival plots showing the association of KRAS expression with overall survival in pancreatic cancer generated using the GEPIA2 database (http://gepia2.cancer-pku.cn). **(b)** Relative mRNA and **(c)** protein expression of KRAS in the MIA PaCa-2 cells cultured under NG or HG conditions. **(d)** Relative mRNA and **(e)** protein expression of KRAS in the spheroids of MIA PaCa-2 cells cultured under NG or HG conditions. **(f)** Immunofluorescence images representing KRAS expression and localization in MIA PaCa-2 cells grown under NG or HG conditions (Scale Bar: 20 µm). **(g)** Immunoblot images representing phospho AKT (pAKT) and total AKT (tAKT) expression in MIA PaCa-2 cells grown under NG or HG. **(h)** Relative lncRNA YIYA expression upon KRAS inhibition (KRASi; Adagrasib dose∼100 nM) in MIA PaCa-2 cells grown under HG conditions. **(i)** Relative lncRNA YIYA expression in HG cultured spheroids of MIA PaCa-2 cells upon KRAS inhibition (KRASi; Adagrasib dose∼100 nM) (The inhibitor was added to the medium mixture containing methylcellulose and HG media, and spheroids were subsequently allowed to form). **(j)** Relative lncRNA YIYA expression upon AKT inhibition (AKTi; PI103, dose∼1 μM) in MIA PaCa-2 cells grown under HG conditions. The glucose concentration used for NG is 5.5 mM, and for HG is 25 mM. Unless otherwise specified, treatments were carried out for 48 h. For quantitative comparisons, bar graphs representing gene expression or protein levels were normalized to the respective housekeeping gene or loading control (either GAPDH or β-actin). Fold change values shown in the graphs are expressed relative to the control, which was set to 1. All results are presented as mean ± SD from at least three independent experiments (n = 3). For comparisons involving multiple groups across varying concentrations and/or time points, two-way ANOVA followed by Tukey’s multiple comparisons post-test was performed. Statistical significance is indicated as (*) p < 0.05, (**) p<0.01, and (***) p < 0.001 versus the control. For pairwise comparisons between two groups, unpaired t-tests were applied.

### High-glucose exposure induces YIYA expression, which in turn drives glycolytic reprogramming consistent with a Warburg phenotype in PDAC cells

Currently, a limited number of studies have explored the potential mechanisms underlying YIYA’s function (17, 27, 37). In this context, a study by Zhen Xing *et al*. (2018) was the first to suggest that YIYA may promote glycolysis in triple-negative breast cancer cells by interacting with cyclin-dependent kinase CDK6 and the glycolytic enzyme fructose bisphosphatase PFK2 (17). Given that YIYA was strongly upregulated alongside the proliferative capacity of PDAC cells in our study, we sought to investigate whether it also influences cellular energy metabolism in PC. To understand the same, we initially examined the metabolic phenotype of MIA PaCa2 cells upon YIYA knockdown. An increased oxygen consumption rate (OCR) was observed under HG, however, siRNA-mediated knockdown of YIYA significantly reduced OCR, indicating that YIYA supports cellular energy metabolism in PC (**Fig. 4a**). Furthermore, total cellular ATP levels were significantly reduced upon siYIYA treatment (**Fig. 4b**). A similar trend was observed in glucose uptake as well-YIYA knockdown caused a substantial reduction in HG-induced increase in expression of glucose transporter-GLUT1 **(Supplementary Fig. 4a and Fig. 4c**), accompanied by a decrease in glucose uptake (**Supplementary Fig. 4b and Fig. 4d**). Overall, these findings suggest that YIYA expression is glucose-dependent, selectively regulates glucose metabolism, and plays a crucial role in glucose uptake through a feedback loop essential for PC growth and survival. Herein, it is well established that following glucose uptake, tumor cells can often adapt a Warburg phenotype—a hallmark of cancer metabolism—where they primarily depend on aerobic glycolysis, converting glucose to lactate even when oxygen is available (6). This metabolic reprogramming is largely supported by key enzymes, such as pyruvate kinase M2 (PKM2) and lactate dehydrogenase (LDH), which enable cells to drive rapid energy production and cellular biosynthesis. In corroboration of the above, in our study, qPCR analysis demonstrated an elevated PKM2 mRNA expression under HG **(Supplementary Fig. 4c**). In addition, immunofluorescence staining also showed an increased PKM2 level and its predominant cytoplasmic localization in HG, and in parallel, immunoblot analysis confirmed an elevated protein expression as well (**Fig. 4e and f**). A similar increase in expression of LDH mRNA levels was also observed under HG in PDAC cells (**Fig. 4g**), along with downregulation of mRNA expression of Mitochondrial Pyruvate Carrier (MPC), which is known to transport pyruvate from the cytosol into the mitochondria, fuelling the TCA cycle (**Fig. 4h**). Importantly, a steep downregulation of PKM2 as well as LDH mRNA levels was observed post YIYA knockdown (**Fig. 4i and Supplementary Fig. 4d**). The changes were also confirmed at the translational level, with immunoblot revealing a significant decrease in PKM2 protein levels upon YIYA knockdown (**Fig. 4j**). These findings indicate YIYA as a key regulator of metabolic proteins, especially PKM2, in PDAC cells and point towards its putative role in promoting glycolysis-driven metabolism under HG conditions. Therefore, to further confirm stimulation of glycolytic flux, we measured extracellular lactate levels in the culture medium, a result of the Warburg effect, as an indicator of lactate secretion. Importantly, lactate levels were elevated in HG **(Supplementary Fig. 4e**), along with mRNA levels of lactate transporter MCT4 **(Supplementary Fig. 4f**). Importantly, YIYA knockdown markedly reduced the lactate levels, which strongly establishes the role of YIYA in maintaining the glycolytic flux (**Fig. 4k**).

**Figure 4:**
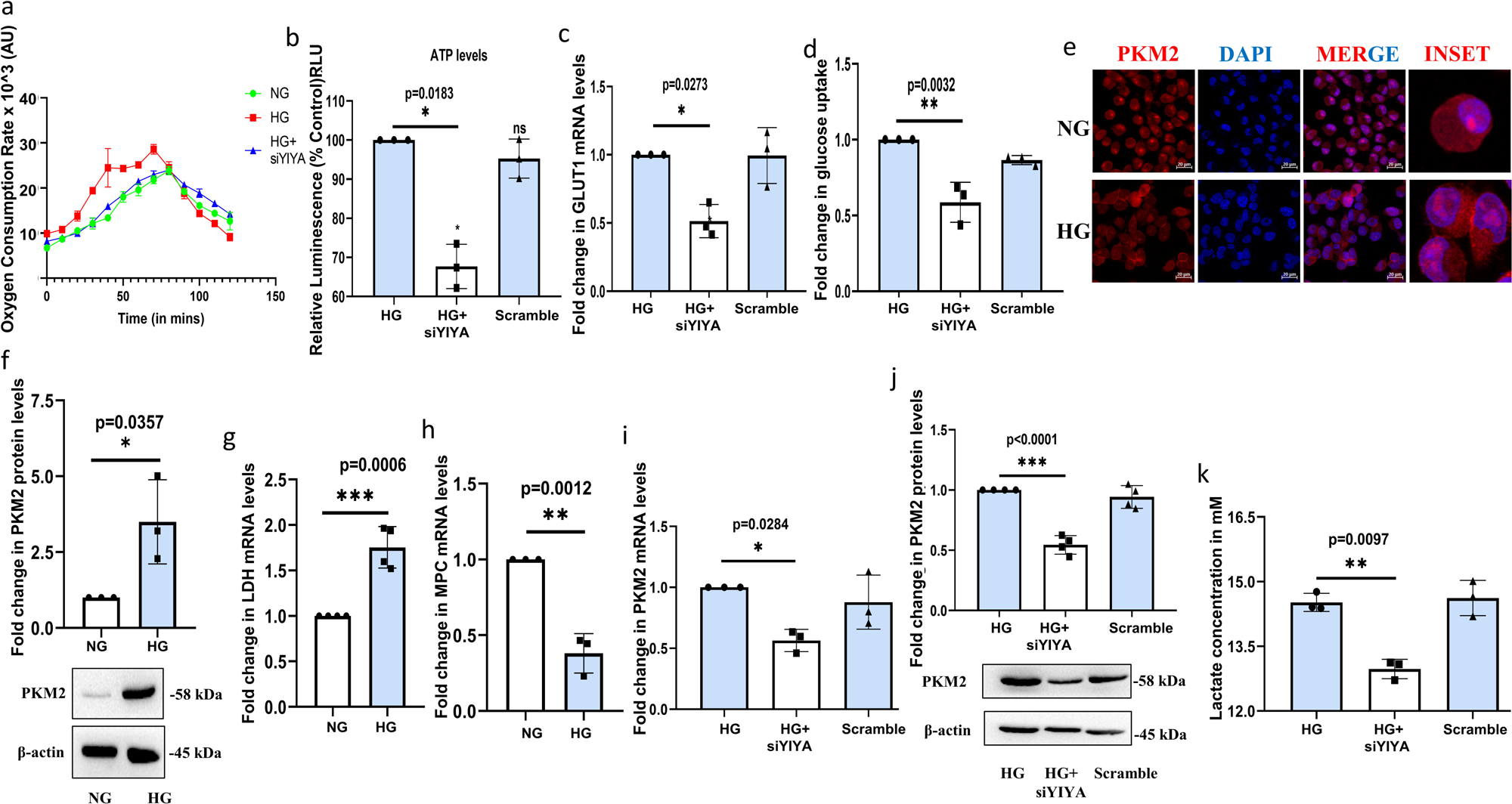
YIYA facilitates a Warburg phenotype under HG conditions. **(a)** Relative change in Oxygen consumption rate (OCR) post-YIYA knockdown in MIA PaCa-2 cells grown under HG conditions. **(b)** Relative change in the ATP levels post-YIYA knockdown in MIA PaCa-2 cells grown under HG conditions. **(c)** Relative mRNA expression of GLUT1 upon YIYA inhibition in MIA PaCa-2 cells grown under HG conditions. **(d)** Relative fold change in glucose uptake post-YIYA inhibition in MIA PaCa-2 cells grown under HG conditions. **(e)** Immunofluorescence images representing PKM2 expression and localization in MIA PaCa-2 cells grown under NG or HG conditions (Scale Bar: 20 µm). **(f)** Immunoblot images representing PKM2 expression in MIA PaCa-2 cells grown under NG or HG. **(g)** Relative mRNA expression of LDH in MIA PaCa-2 cells grown under NG or HG. **(h)** Relative mRNA expression of MPC in MIA PaCa-2 cells grown under NG or HG. **(i)** Relative mRNA expression of PKM2 in HG cultured MIA PaCa-2 cells upon YIYA inhibition. **(j)** Immunoblot images representing PKM2 expression in MIA PaCa-2 cells grown under HG upon YIYA inhibition. **(k)** Relative fold change in extracellular lactate concentration in HG cultured MIA PaCa-2 cells upon YIYA inhibition. The glucose concentration used for NG is 5.5 mM, and for HG is 25 mM. The final concentration of siRNA used for transfection was 40 nM. Unless otherwise specified, treatments were carried out for 48 h. For quantitative comparisons, bar graphs representing gene expression or protein levels were normalized to the respective housekeeping gene or loading control (either GAPDH or β-actin). Fold change values shown in the graphs are expressed relative to the control, which was set to 1. All results are presented as mean ± SD from at least three independent experiments (n = 3). For comparisons involving multiple groups across varying concentrations and/or time points, two-way ANOVA followed by Tukey’s multiple comparisons post-test was performed. Statistical significance is indicated as (*) p < 0.05, (**) p<0.01, and (***) p < 0.001 versus the control. For pairwise comparisons between two groups, unpaired t-tests were applied.

### Through direct interactions with KRAS and PKM2, YIYA influences protein turnover by modulating autophagy, thereby impacting metabolic homeostasis in PDAC cells

We next aimed to investigate the molecular mechanism by which YIYA regulates the specific cellular metabolic state upon exposure to HG. Interestingly, KRAS mRNA levels remained largely unchanged (**Fig. 5a**) after YIYA knockdown; while we observed a marked reduction in KRAS protein levels (**Fig. 5b**), suggesting that YIYA regulates KRAS post-transcriptionally. Building on our previous findings that hyperglycemia enhances KRAS signalling, we hypothesized that the lncRNA YIYA may contribute to this process through interactions with key oncogenic or metabolic proteins. Specifically, given YIYA’s observed regulatory role and the importance of mutant KRAS in PDAC, we tested whether YIYA physically associates with KRAS under hyperglycaemic conditions. RNA immunoprecipitation (RIP) using a KRAS-specific antibody revealed strong enrichment of YIYA lncRNA in KRAS-bound complexes in HG-treated MIA PaCa-2 cells, indicating that YIYA physically interacts with KRAS (**Fig. 5c**). Given that YIYA interacts with KRAS and that siYIYA reduces the protein levels of both KRAS and PKM2, we speculated that KRAS and PKM2 may exist in a functional complex, and that YIYA might facilitate or stabilize this interaction. Although KRAS is classically known for its role in signalling pathways, its non-canonical roles are also reported (38). To explore this potential crosstalk, we initially treated PC cells with a KRAS inhibitor (Adagrasib) and observed that this led to a marked reduction in PKM2 mRNA and protein expression (**Fig. 5d and e**), suggesting that PKM2 levels are maintained *via* KRAS signalling. Subsequently, we performed a co-immunoprecipitation (Co-IP) assay using a KRAS antibody and probed for PKM2. We observed a robust interaction between KRAS and PKM2, which suggests that these two proteins may form a complex under HG conditions (**Fig. 5f**). To determine whether YIYA could be part of the complex, we next performed RIP using a PKM2 antibody, which revealed a strong enrichment of YIYA RNA with PKM2 as well (**Fig. 5g**). Together, these results raise the possibility that YIYA connects oncogenic KRAS signalling and glycolytic reprogramming by acting as a scaffold for the KRAS-PKM2 complex in HG-treated PDAC cells. To understand how YIYA influences the KRAS protein levels, we next explored whether their reduction upon siYIYA could be due to enhanced protein degradation. Since autophagy and proteasomal degradation pathways are common routes for protein turnover (39,40), we treated siYIYA-transfected cells with chloroquine (CQ), an autophagosome-lysosome fusion inhibitor that blocks autophagic flux. We found that KRAS protein levels were partially restored in the siYIYA plus CQ condition (**Fig. 5h**), suggesting that in the absence of YIYA, KRAS protein is destabilized and degraded *via* autophagy. However, proteasomal inhibition with MG132 showed no significant accumulation of KRAS (data not included in results). Thus, YIYA may protect KRAS from degradation, ensuring the persistence of oncogenic and glycolytic signalling under hyperglycemic stress.

**Figure 5:**
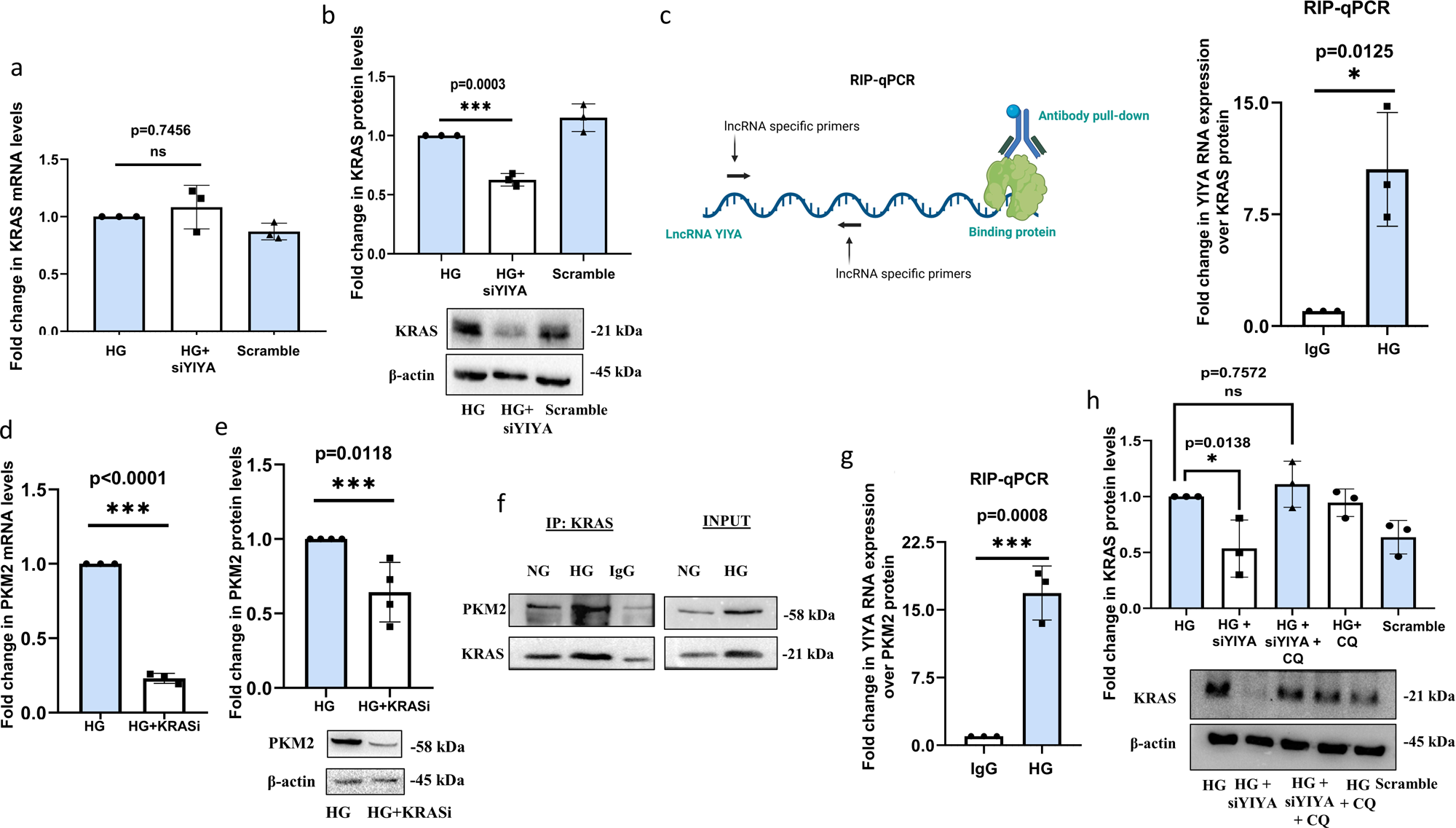
YIYA binds to KRAS and modulates KRAS–PKM2 complex formation and stability. **(a)** Relative mRNA and **(b)** protein expression of KRAS upon YIYA inhibition in MIA PaCa-2 cells grown under HG conditions. **(c)** Interaction of lncRNA YIYA with KRAS protein in MIA PaCa-2 cells grown under HG conditions, as analyzed using RIP-qPCR (KRAS was immunoprecipitated using a specific antibody, and the associated RNA was analyzed by qPCR). IgG served as a control. RIP-qPCR illustration prepared with Biorender software. **(d)** Change in mRNA expression of PKM2 upon KRAS inhibition (KRASi; Adagrasib dose∼100 nM) in MIA PaCa-2 cells grown under HG conditions. **(e)** Immunoblot images representing PKM2 protein expression in MIA PaCa-2 cells grown under HG upon KRAS inhibition (Adagrasib dose∼100 nM). **(f)** Interaction of PKM2 protein with KRAS protein in MIA PaCa-2 cells grown under NG or HG conditions, as analyzed using CO-IP (KRAS was immunoprecipitated using a specific antibody, and the precipitated complexes were analyzed by SDS–PAGE followed by immunoblotting with a PKM2 antibody.) **(g)** Interaction of lncRNA YIYA with PKM2 protein in MIA PaCa-2 cells grown under HG conditions, as analyzed using RIP-qPCR (PKM2 was immunoprecipitated using a specific antibody, and the associated RNA was analyzed by qPCR). IgG served as a control. **(h)** Immunoblot images representing KRAS protein expression in MIA PaCa-2 cells grown under HG upon YIYA inhibition and autophagy inhibition (CQ∼10 μM). The glucose concentration used for NG is 5.5 mM, and for HG is 25 mM. The final concentration of siRNA used for transfection was 40 nM. Unless otherwise specified, treatments were carried out for 48 h. For quantitative comparisons, bar graphs representing gene expression or protein levels were normalized to the respective housekeeping gene or loading control (either GAPDH or β-actin). Fold change values shown in the graphs are expressed relative to the control, which was set to 1. All results are presented as mean ± SD from at least three independent experiments (n = 3). For comparisons involving multiple groups across varying concentrations and/or time points, two-way ANOVA followed by Tukey’s multiple comparisons post-test was performed. Statistical significance is indicated as (*) p < 0.05, (**) p<0.01, and (***) p < 0.001 versus the control. For pairwise comparisons between two groups, unpaired t-tests were applied.

## Discussion

Pancreatic ductal adenocarcinoma (PDAC) is a formidable malignancy characterised by late diagnosis, chemoresistance, and a largely altered metabolic and epigenetic landscape. Many PDAC patients present with hyperglycemia or new-onset diabetes. This raises a question about the causal relationship between dysregulated glucose metabolism and PDAC progression (41). Indeed, hyperglycemia not only supplies energy to rapidly dividing tumor cells but also acts as a signalling molecule that regulates cellular transcriptional and epigenetic changes (42). However, the molecular mediators that translate hyperglycaemic stress into tumor-promoting phenotypes remain inadequately defined.

Our study adds to the growing body of evidence linking HG exposure to PDAC progression by identifying lncRNA YIYA as a key regulatory node in this axis. Unlike traditional oncogenes or metabolic enzymes, lncRNAs provide a unique layer of regulation through their modular functions, acting as scaffolds, decoys, or guides, often within specific physiological contexts (42). We demonstrate that YIYA is not only responsive to hyperglycaemic conditions but also coordinates key oncogenic and metabolic players like KRAS and PKM2, thus offering a mechanistic link between glucose-induced signalling and tumour metabolism. One of the most striking observations from our study is the ability of YIYA to stabilize the KRAS protein under hyperglycaemic conditions. This highlights a post-transcriptional layer of control exerted by lncRNAs, potentially by shielding protein complexes from protein degradation processes like autophagy. In this context, our findings also align with recent reports implicating non-canonical functions of KRAS beyond its classical signalling roles, including protein–protein interactions that modulate metabolic plasticity (43,44). The formation of a YIYA–KRAS–PKM2 complex not only provides novel insight into metabolic reprogramming but also opens avenues for therapeutic targeting of such tripartite regulatory axes. It is also important to note that YIYA interacts not only with a master oncogenic regulator such as KRAS but also with a key metabolic regulator, PKM2. In PDAC, PKM2 is known to contribute to tumour progression through its canonical role in driving aerobic glycolysis (the Warburg effect), as well as through noncanonical functions, including acting as a protein kinase, translocating to the nucleus to influence transcriptional programs, and interacting with oncogenic signalling pathways (45). These multifaceted activities establish PKM2 as a central metabolic and regulatory hub in PC.

It is important to note that KRAS mutations are nearly ubiquitous in PC, with the most common variants being G12D, G12V, and G12R, while G12C occurs at a lower frequency, and they have attracted major therapeutic interest (46). These mutations drive constitutive KRAS activation, which sustains oncogenic signalling. Clinically, the presence of specific KRAS alleles can have an impact on disease prognosis and therapeutic outcomes (30). For instance, G12D-mutant tumors are often associated with aggressive biology and poor survival (47), while G12C mutations have become a focus of drug development (46). Small-molecule KRAS G12C inhibitors such as adagrasib have shown clinical efficacy in lung and colorectal cancers, and early-phase clinical trials are currently running for PDAC (35). Recent studies suggest that KRAS not only drives canonical oncogenic signalling but also interacts with non-coding RNAs in PDAC. Several lncRNAs, like MALAT1 and NUTF2P3-01 (48,49), have been reported to modulate KRAS expression or signalling output, either by stabilizing KRAS mRNA, regulating its upstream activators, or scaffolding downstream effectors. Herein, our findings that YIYA stabilizes KRAS protein and promotes KRAS–PKM2 complex formation extend this pattern by highlighting an lncRNA-mediated mechanism of post-translational regulation, which emphasizes the emerging role of lncRNAs as modulators of KRAS oncogenic networks in PDAC.

Furthermore, from a metabolic perspective, our data reinforce the relevance of the Warburg effect in hyperglycemia-exposed PDAC cells. Enhanced aerobic glycolysis, lactate production, and downregulation of mitochondrial transporters like MPC support the idea that hyperglycemia potentiates a glycolytic switch, and YIYA appears to be a necessary modulator of this phenotype. Several other lncRNAs, such as PVT1, MALAT1, and HOTAIR, have also been linked to glucose metabolism in PDAC (50–52), further suggesting that non-coding RNAs might act as metabolic rheostats in cancer. However, YIYA’s ability to simultaneously interface with metabolic enzymes and oncogenic drivers makes it uniquely positioned in the PDAC’s adaptive landscape. Our findings also prompt important questions about the epigenetic consequences of metabolic stress. Previous studies, including work from our lab on the lncRNA HULC (22), have shown that high glucose exposure can alter histone marks and chromatin accessibility. In the current study, ChIP-qPCR data indicate that YIYA modulates epigenetic marks at the promoters of proliferation-associated genes, albeit indirectly. The cytoplasmic localization of YIYA suggests it may influence the epigenome through secondary regulatory cascades, perhaps involving signalling intermediates or chromatin modifiers responsive to metabolic flux. Future studies can explore whether YIYA alters α-ketoglutarate–dependent histone demethylases or participates in feedback loops involving histone acetylation and DNA methylation.

Although our work is the first to characterize YIYA in the context of PDAC, there are only limited reports on this lncRNA in other cancers. For example, studies in breast cancer have shown that YIYA promotes cell cycle progression by interacting with cyclin-dependent kinases and modulating glycolytic metabolism, while in colon cancer, YIYA has been linked to tumor cell proliferation via EZH2 and NKD2 (27). These findings, together with our data, suggest that YIYA functions as a context-dependent oncogenic lncRNA, with the capacity to interface with both metabolic and signalling pathways across tumor types. This broader oncogenic role strengthens the rationale for investigating YIYA as a potential therapeutic target beyond PDAC. The clinical implications of our study are twofold. First, given the high prevalence of diabetes or impaired glucose tolerance in PDAC patients, monitoring lncRNA expression, especially glucose-sensitive ones like YIYA, could serve as a biomarker for disease aggressiveness or treatment response. Second, the fact that YIYA appears to protect key oncogenic proteins from degradation suggests a potential vulnerability. Inhibiting YIYA may destabilize oncogenic signalling networks and re-sensitize tumours to chemotherapy. Our findings set the stage for multiple future directions. One avenue involves mapping the YIYA interactome under different metabolic conditions to identify cofactors involved in KRAS–PKM2 stabilization. Another critical direction is to explore how intermittent hyperglycemic spikes, often seen in diabetic patients, alter YIYA expression and function over time. Furthermore, the *in vivo* relevance of our study in diabetic mouse models of PDAC will be essential to validate the therapeutic potential of targeting this axis. In conclusion, this study positions lncRNA YIYA as a novel metabolic-epigenetic integrator in hyperglycemia-driven pancreatic cancer. By connecting glucose sensing, oncogene stabilization, and metabolic rewiring, YIYA emerges as a promising biomarker and therapeutic target in malignancies that desperately need both.

## ASSOCIATED CONTENT

### Data Availability Statement

The data underlying this study are available in the published article and its Supporting Information.

## SUPPORTING INFORMATION

### NOTES

The authors declare that they have no conflict of interest.

## ACKNOWLEDGMENTS

This research work was funded by two projects of the Department of Science and Technology, Government of India, New Delhi, under the SERB-ANRF (EMR/2016/006788) (CRG/2022/003241). The authors are grateful to Birla Institute of Technology and Science, Pilani, for providing lab facilities and infrastructure.

---

**Supplementary Figure 1: (a)** Immunofluorescence images representing PCNA expression in MIA PaCa-2 cells grown under NG or switched from NG to HG conditions (Scale Bar: 20 µm). **(b)** Immunofluorescence images representing PCNA expression in MIA PaCa-2 cells grown under HG or switched from HG to NG conditions (Scale Bar: 20 µm). **(c)** Phase-contrast images of spheroids of MIA PaCa-2 cells grown under NG or HG conditions (Scale Bar: 100 µm). **(d)** Relative lncRNA YIYA expression of MIA PaCa-2 cells cultured under NG or switched to NG to HG conditions. **(e)** Relative lncRNA YIYA expression of MIA PaCa-2 cells cultured under HG or switched to HG to NG conditions. The glucose concentration used for NG is 5.5 mM, and for HG is 25 mM. For quantitative comparisons, bar graphs representing gene expression or protein levels were normalized to the respective housekeeping gene or loading control (either GAPDH or β-actin). Fold change values shown in the graphs are expressed relative to the control, which was set to 1. All results are presented as mean ± SD from at least three independent experiments (n = 3). For comparisons involving multiple groups across varying concentrations and/or time points, two-way ANOVA followed by Tukey’s multiple comparisons post-test was performed. Statistical significance is indicated as (*) p < 0.05, (**) p<0.01, and (***) p < 0.001 versus the control. For pairwise comparisons between two groups, unpaired t-tests were applied.

**Supplementary Figure 2: (a)** Phase-contrast micrographs showing morphological changes in MIA PaCa-2 cells after siRNA-mediated silencing of lncRNA YIYA under HG conditions. (Scale bar: 50 µm). **(b)** Relative mRNA expression of Caspase-3 in MIA PaCa2 cells cultured under HG conditions and upon YIYA knockdown. **(c)** Wound healing assay of MIA PaCa-2 cells following incubation under HG conditions or treatment with siYIYA. Images were captured at 0 h and 48 h. (Scale bar: 100 µm). **(d)** Kaplan–Meier survival analysis obtained from the lnc2cancer database (http://www.bio-bigdata.com/lnc2cancer), illustrating the relationship between lncRNA YIYA expression levels and patient survival in pancreatic cancer. The glucose concentration used for NG is 5.5 mM, and for HG is 25 mM. The final concentration of siRNA used for transfection was 40 nM. Unless otherwise specified, treatments were carried out for 48 h. For quantitative comparisons, bar graphs representing gene expression or protein levels were normalized to the respective housekeeping gene or loading control (either GAPDH or β-actin). Fold change values shown in the graphs are expressed relative to the control, which was set to 1. All results are presented as mean ± SD from at least three independent experiments (n = 3). For comparisons involving multiple groups across varying concentrations and/or time points, two-way ANOVA followed by Tukey’s multiple comparisons post-test was performed. Statistical significance is indicated as (*) p < 0.05, (**) p<0.01, and (***) p < 0.001 versus the control. For pairwise comparisons between two groups, unpaired t-tests were applied.

**Supplementary Figure 3: (a)** Immunoblot images representing phospho ERK (pERK) and total ERK (tERK) expression post KRAS inhibition (KRASi; Adagrasib∼100 nM) in MIA PaCa-2 cells grown under HG. **(b)** Immunoblot images representing KRAS expression post KRAS inhibition (KRASi; Adagrasib∼100 nM) in MIA PaCa-2 cells grown under HG. **(c)** Phase-contrast images of spheroids of MIA PaCa-2 cells grown under HG or post KRAS inhibition (KRASi; Adagrasib∼100 nM) grown under HG conditions, from day 1 to day 7 (Scale Bar: 100 µm). **(d)** Change in area of spheroids of MIA PaCa-2 cells grown under HG or post KRAS inhibition (KRASi; Adagrasib∼100 nM) grown under HG conditions, from day 1 to day 7. **(e)** Immunoblot images representing phospho AKT (pAKT) and total AKT (tAKT) expression post AKT inhibition (AKTi; PI103∼1 µM) in MIA PaCa-2 cells grown under HG. The glucose concentration used for NG is 5.5 mM, and for HG is 25 mM. Unless otherwise specified, treatments were carried out for 48 h. For quantitative comparisons, bar graphs representing gene expression or protein levels were normalized to the respective housekeeping gene or loading control (either GAPDH or β-actin). Fold change values shown in the graphs are expressed relative to the control, which was set to 1. All results are presented as mean ± SD from at least three independent experiments (n = 3). For comparisons involving multiple groups across varying concentrations and/or time points, two-way ANOVA followed by Tukey’s multiple comparisons post-test was performed. Statistical significance is indicated as (*) p < 0.05, (**) p<0.01, and (***) p < 0.001 versus the control. For pairwise comparisons between two groups, unpaired t-tests were applied.

**Supplementary Figure 4: (a)** Relative mRNA expression of GLUT1 in MIA PaCa-2 cells grown under NG or HG. **(b)** Relative fold change in glucose uptake in MIA PaCa-2 cells grown under NG or HG. **(c)** Relative mRNA expression of PKM2 in MIA PaCa-2 cells grown under NG or HG. **(d)** Relative mRNA expression of LDH in HG cultured MIA PaCa-2 cells upon YIYA inhibition. **(e)** Relative extracellular lactate concentration in MIA PaCa-2 cells grown under NG or HG. **(f)** Relative mRNA expression of MCT4 in MIA PaCa-2 cells grown under NG or HG. The glucose concentration used for NG is 5.5 mM, and for HG is 25 mM. The final concentration of siRNA used for transfection was 40 nM. Unless otherwise specified, treatments were carried out for 48 h. For quantitative comparisons, bar graphs representing gene expression or protein levels were normalized to the respective housekeeping gene or loading control (either GAPDH or β-actin). Fold change values shown in the graphs are expressed relative to the control, which was set to 1. All results are presented as mean ± SD from at least three independent experiments (n = 3). For comparisons involving multiple groups across varying concentrations and/or time points, two-way ANOVA followed by Tukey’s multiple comparisons post-test was performed. Statistical significance is indicated as (*) p < 0.05, (**) p<0.01, and (***) p < 0.001 versus the control. For pairwise comparisons between two groups, unpaired t-tests were applied.

