## Supplementary figures and images for "LncRNA YIYA enhances pancreatic cancer proliferation under high-glucose conditions through RAS–PKM2–mediated metabolic reprogramming that reinforces the Warburg phenotype"

### Supplementary Figures

#### Supplementary Figure 1

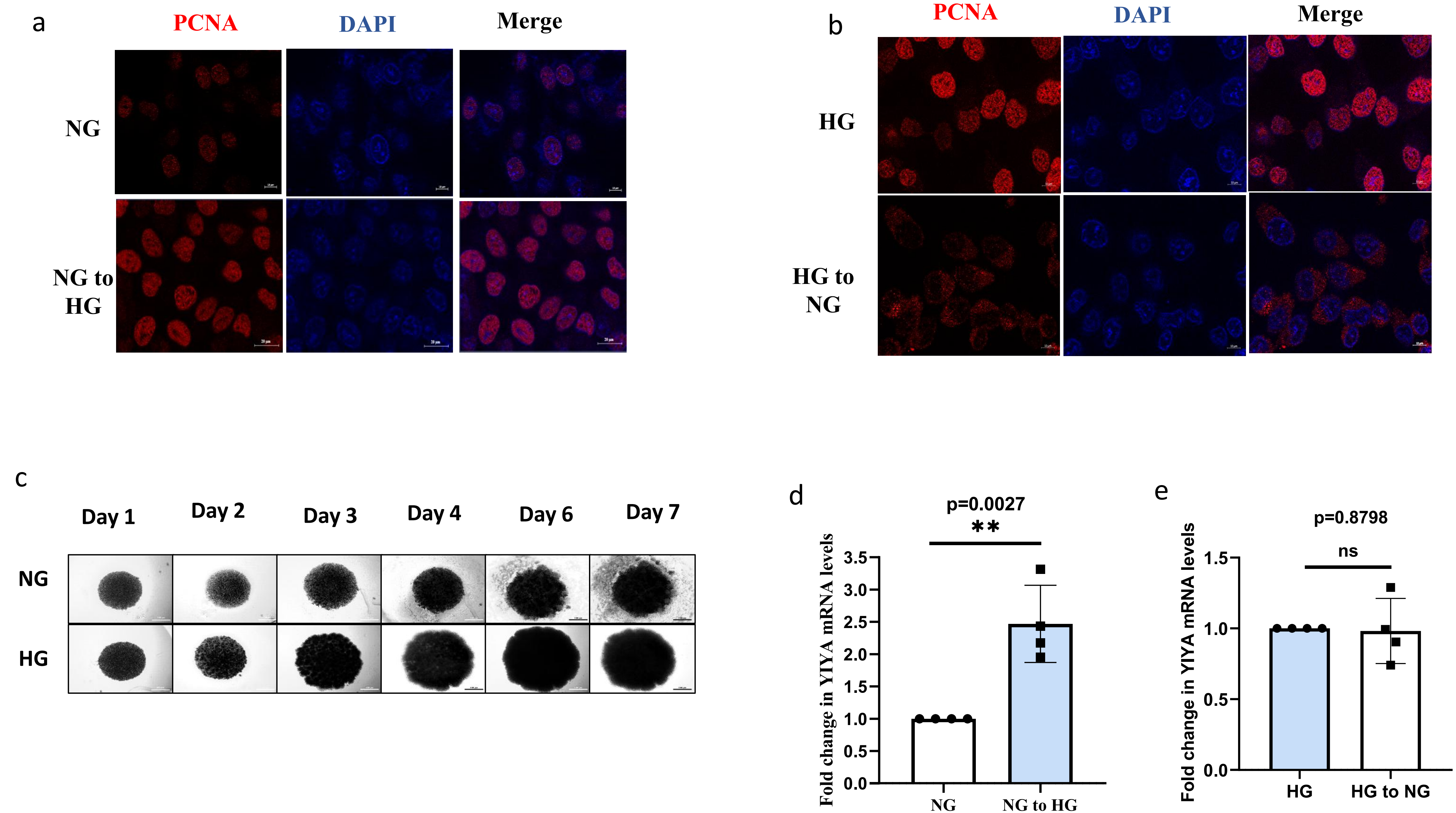

#### Supplementary Figure 2

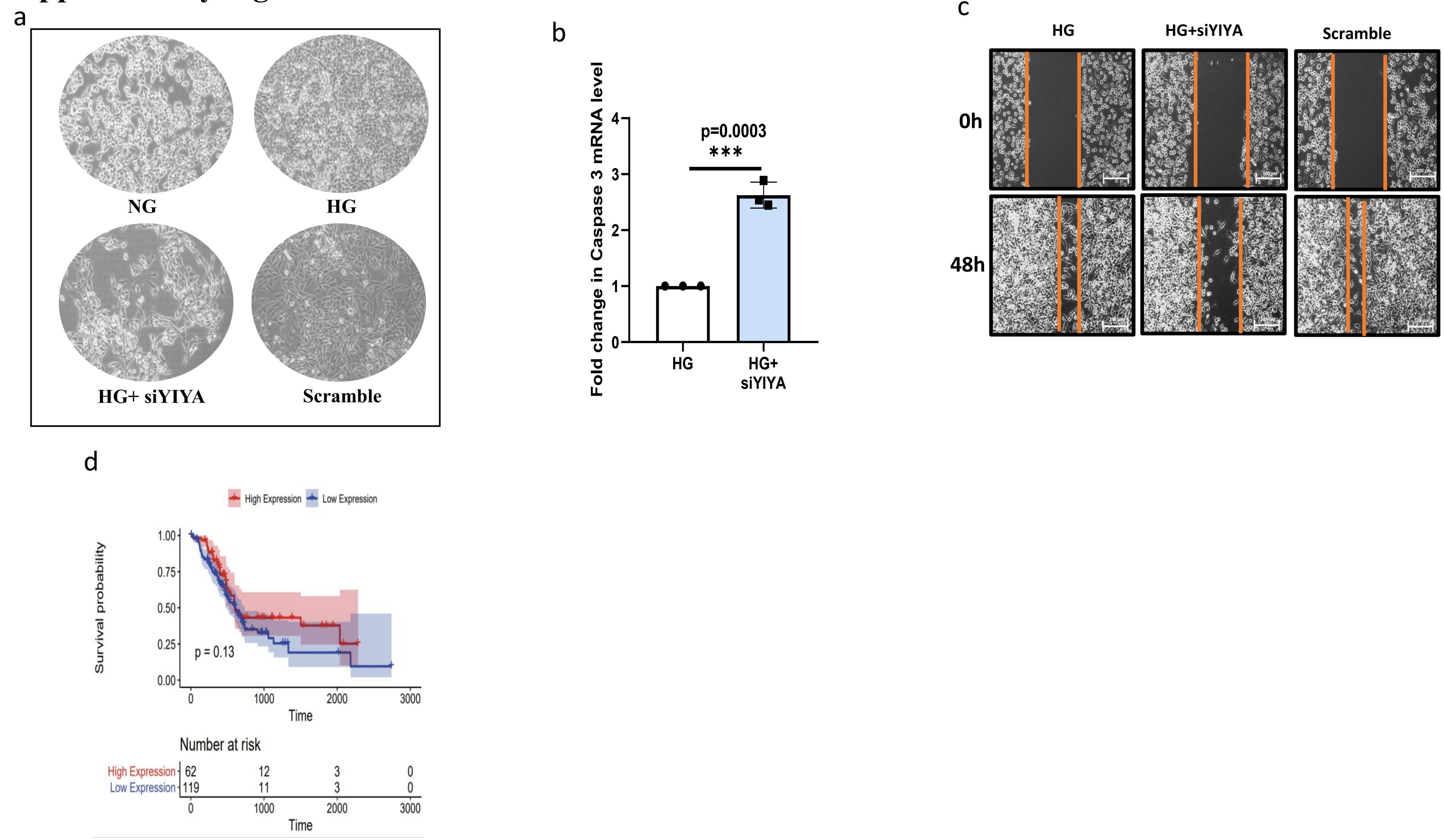

#### Supplementary Figure 3

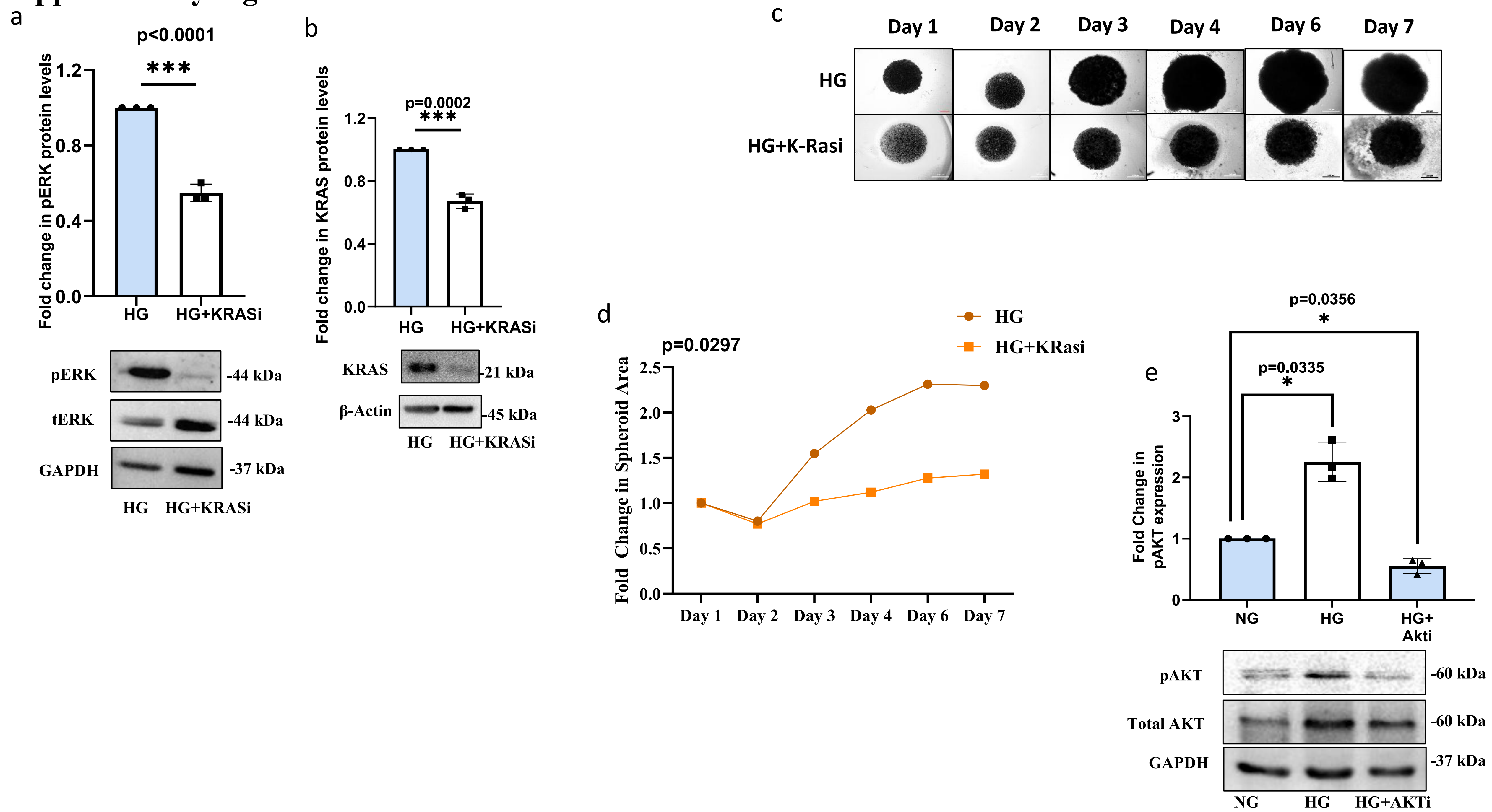

#### Supplementary Figure 4

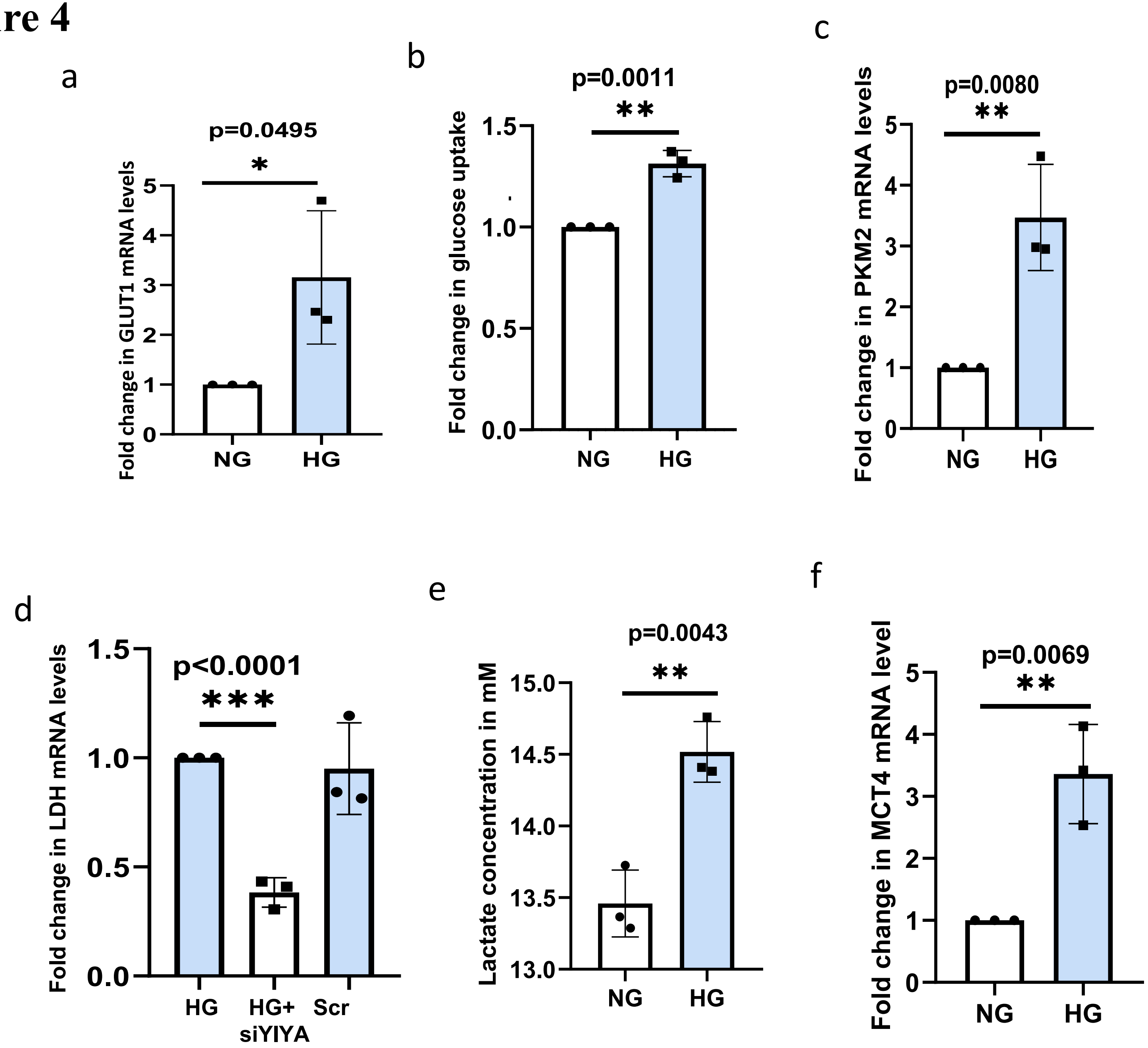
